# AGO1 and HSP90 buffer different genetic variants in *Arabidopsis thaliana*

**DOI:** 10.1101/2022.06.27.497824

**Authors:** Tzitziki Lemus, G. Alex Mason, Kerry L. Bubb, Cristina M. Alexandre, Christine Queitsch, Josh T. Cuperus

## Abstract

Argonaute 1 (AGO1), the principal protein component of microRNA-mediated regulation, plays a key role in plant growth and development. AGO1 physically interacts with the chaperone HSP90, which buffers cryptic genetic variation in plants and animals. We sought to determine whether genetic perturbation of *AGO1* in *Arabidopsis thaliana* would also reveal cryptic genetic variation, and if so, whether *AGO1*-dependent loci overlap with those dependent on HSP90. To address these questions, we introgressed a hypomorphic mutant allele of *AGO1* into a set of mapping lines derived from the commonly used *Arabidopsis* strains Col-0 and L*er*. Although we identified several cases in which AGO1 buffered genetic variation, none of the *AGO1*-dependent loci overlapped with those buffered by HSP90 for the traits assayed. We focused on one buffered locus where *AGO1* perturbation uncoupled the traits days to flowering and rosette leaf number, which are otherwise closely correlated. Using a bulk segregant approach, we identified a non-functional L*er hua2* mutant allele as the causal AGO1-buffered polymorphism. Introduction of a non-functional *hua2* allele into a Col-0 *ago1* mutant background recapitulated the L*er-*dependent *ago1* phenotype, implying that coupling of these traits involves different molecular players in these closely related strains. Taken together, our findings demonstrate that even though AGO1 and HSP90 buffer genetic variation in the same traits, these robustness regulators interact epistatically with different genetic loci, suggesting that higher-order epistasis is uncommon.

**Article Summary:** Argonaute 1 (AGO1), a key player in plant development, interacts with the chaperone HSP90 which buffers environmental and genetic variation. We found that *AGO1* buffers environmental and genetic variation in the same traits; however, AGO1-dependent and HSP90-dependent loci do not overlap. Detailed analysis of a buffered locus found that a non-functional *HUA2* allele decouples days to flowering and rosette leaf number in an AGO1-dependent manner, suggesting the AGO1-dependent buffering acts at the network level.

## Introduction

Genetic networks rely on various types of feedback loops, redundancy, and other mechanisms like chaperones and small RNAs to ensure phenotypic robustness in spite of environmental or genetic perturbations (Rutherford and Lindquist 1998; Queitsch *et al*. 2002; Masel and Siegal 2009; Whitacre 2012; Lempe *et al*. 2013; Lachowiec *et al*. 2018; Zabinsky *et al*. 2019). Network disruptions decrease environmental and developmental robustness and, dependent on their nature, increase phenotypic variation in a trait or affect organismal phenotypes more broadly. For example, perturbation of the essential chaperone HSP90 broadly increases phenotypic variation in plants, fungi, and animals, with many organismal traits affected in a background-specific manner (Rutherford and Lindquist 1998; Queitsch *et al*. 2002; Yeyati *et al*. 2007; Sangster *et al*. 2008b; a; Jarosz and Lindquist 2010; Rohner *et al*. 2013; Karras *et al*. 2017; Zabinsky *et al*. 2019). When fully functional, HSP90 chaperones a select but highly diverse group of client proteins, including many kinases, receptors and transcription factors with crucial roles in development (Schopf *et al*. 2017). When chaperone function is perturbed, client proteins encoding genetic variants may fail to mature or fold differently, leading to pathway failure or rewiring (Dorrity *et al*. 2018) and hence altered phenotypes (Zabinsky *et al*. 2019). The phenomenon that HSP90 keeps genetic variation phenotypically silent and HSP90 perturbation allows its expression has become known as phenotypic capacitance (Rutherford and Lindquist 1998; Masel and Siegal 2009) – a different term for epistasis (Zabinsky *et al*. 2019). In contrast to the traditional definition of epistasis, which describes the non-reciprocal interaction of two loci, phenotypic capacitance is an epistasis phenomenon in which one locus, *e*.*g*., *HSP90*, interacts with several others. HSP90 perturbation can increase phenotypic variation even in the absence of genetic variation, presumably because subtle differences in microenvironment, developmental stage, or cell state lead to inhibition of different client proteins among individuals in a seemingly stochastic manner (Zabinsky *et al*. 2019)..

Another important source of developmental and environmental robustness is post-transcriptional regulation by small RNAs. Small RNAs regulate the expression of their target genes in a sequence-specific manner. In plants, most endogenous post-transcriptional gene regulation is mediated by AGO1 loaded with microRNAs (miRNAs, MIR) (Axtell 2013; Bologna and Voinnet 2014). In animals, some microRNAs are known to buffer stochastic (Hilgers *et al*. 2010), environmental (Li *et al*. 2009), and genetic variation (Cassidy *et al*. 2013). MicroRNAs play major roles throughout plant development, including in the onset of flowering, an irreversible developmental transition of outsized effect on reproductive success in annual plants (Dong *et al*. 2022). In particular, the MIR156 and MIR172 gene families are essential for fine-tuning expression of the complex gene network that determines the number of days until flowering is initiated and the number of rosette leaves at this developmental stage. Misregulation or mutation of their gene targets alters both traits in the crucifer model *Arabidopsis thaliana* (Aukerman and Sakai 2003; Yamaguchi *et al*. 2009; Wu *et al*. 2009).

In *Arabidopsis*, the traits days to flowering (*i*.*e*., flowering time, onset of flowering) and rosette leaf number are so closely linked that the traits are often used interchangeably. This close correlation reflects the need for sufficient vegetative tissue (*i*.*e*., rosette leaves) to produce the resources for flowering and seed development. Because of the irreversible nature of the transition from the vegetative to the reproductive stage in *Arabidopsis*, the coupling of these traits is crucial for reproductive success. *Arabidopsis* mutants that flower with as few as three or four adult leaves develop very few seeds and often show weakened growth habits. Uncoupling of flowering time and rosette leaf number occurs in some early and late flowering time mutants (Pouteau *et al*. 2004) and in response to treatment with nitrogen dioxide (Takahashi and Morikawa 2014); however, the mechanistic underpinnings for this uncoupling remain unknown. Studies in several organisms suggest that AGO proteins are chaperoned by HSP90. HSP90 physically interacts with AGO proteins in yeast (Wang *et al*. 2013; Okazaki *et al*. 2018), flies (Iwasaki *et al*. 2010; Miyoshi *et al*. 2010; Gangaraju *et al*. 2011), humans (Johnston *et al*. 2010; Gangaraju *et al*. 2011), *Tetrahymena* (Woehrer *et al*. 2015), and plants (Iki *et al*. 2010, 2012). Because miRNAs buffer environmental and genetic perturbations and AGO1 interacts with HSP90, we set out to investigate the extent to which *AGO1* perturbation affects phenotypic variation in isogenic *Arabidopsis* seedlings and buffers genetic variation in divergent backgrounds, and *AGO1*-dependent loci overlap with HSP90-dependent loci. We find that *AGO1* perturbation can significantly increase phenotypic variation in morphological and quantitative traits in isogenic seedlings. *AGO1* perturbation also buffers the phenotypic effects of genetic variation between two divergent backgrounds. However, none of the *AGO1*-buffered loci overlapped with those buffered by HSP90, consistent with a prevalence of first-order epistatic interactions relative to higher-order epistasis. Lastly, our detailed investigation of one such buffered locus reveals that the coupling of the fitness-relevant traits days to flowering and rosette leaf number relies on different molecular players in these commonly used strains of *Arabidopsis*.

## Results

### Genetic perturbation of *AGO1* increases phenotypic variation in isogenic *Arabidopsis* seedlings

To determine if perturbation of *AGO1* leads to increased phenotypic variation in isogenic seedlings, we examined several morphological and quantitative traits of two hypomorphic *ago1* mutants, *ago1-46* (Smith *et al*. 2009), and *ago1-27* (Morel 2002), the former being a less severe mutant than the latter. Ten-day old isogenic seedlings of *ago1-46* and *ago1-27* showed increased phenotypic variation in morphological traits such as lesions in cotyledons (Mason *et al*. 2016), rosette symmetry, and organ defects compared to isogenic wild-type seedlings (**Figure 1, Supplemental Table 1**). As expected, the more severe *ago1-27* mutant showed more abnormal phenotypes than the less severe *ago1-46* mutant. Next, we examined hypocotyl length in the dark, a quantitative trait that shows increased variation in response to HSP90 perturbation (Queitsch *et al*. 2002; Sangster *et al*. 2008b). Similar to our previous results (Queitsch *et al*. 2002; Sangster *et al*. 2008b), *ago1-27* dark-grown seedlings showed a different mean value (p < 2.3e^−16^, Wilcoxon test) and significantly greater variance of hypocotyl length than wild-type seedlings (p = 0.0002, Levene’s test) (**Figure 1B, Supplemental Table 1**). The less severe *ago1-46* seedlings also showed a different mean value (p-value = 3.6e^−05^, Wilcoxon test) and greater variance of hypocotyl length compared to wild-type seedlings (p-value = 0.00004, Levene’s test). Based on these results, AGO1 maintains phenotypic robustness and buffers developmental noise among isogenic seedlings in a manner similar to HSP90.

**Figure 1.**
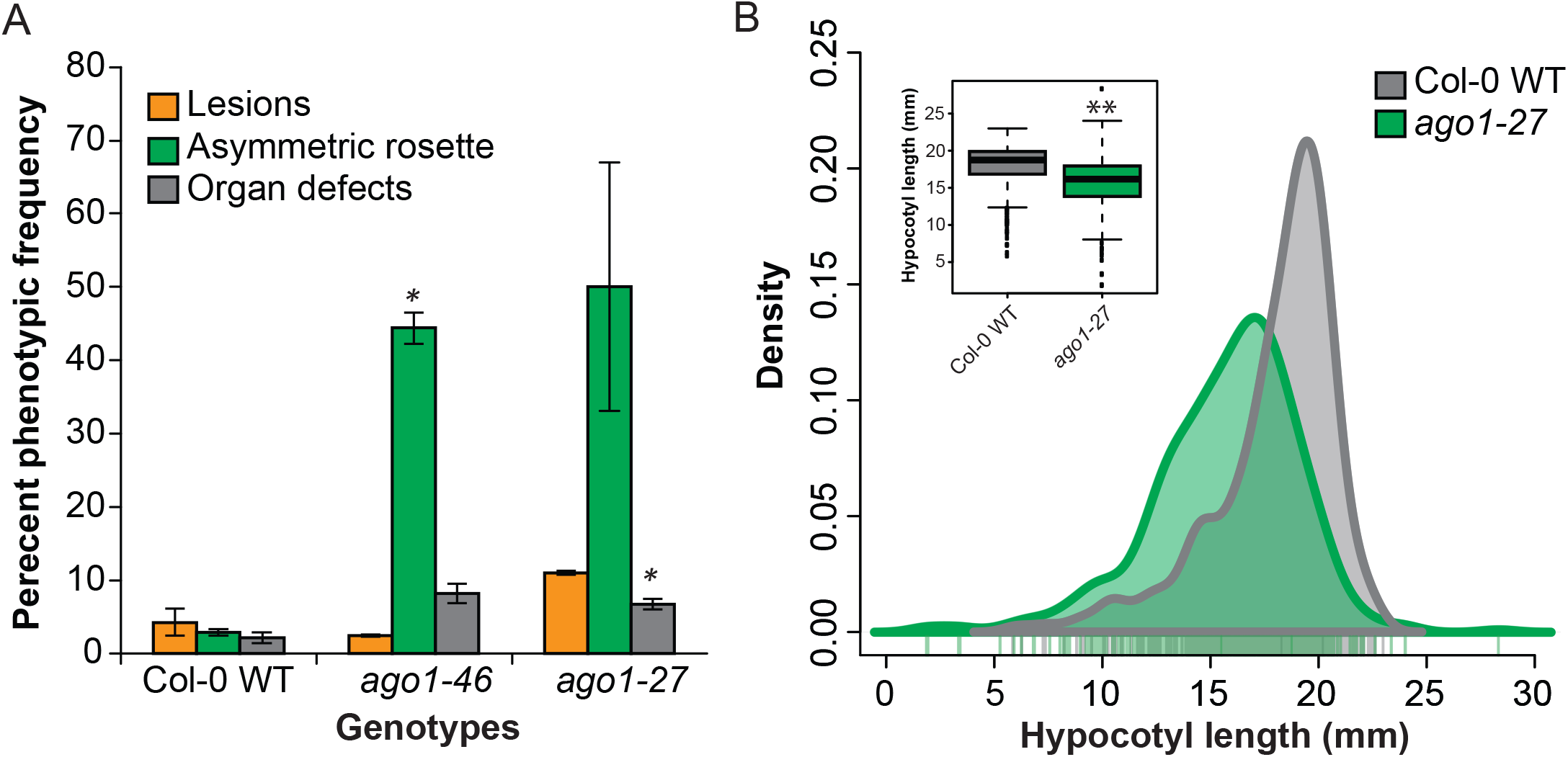
Perturbation of AGO1 increases phenotypic variation among isogenic seedlings. **A**) Early seedling trait measures for wild-type (Col-0 WT), *ago1-46*, and *ago1-27* seedlings. Ten-day-old seedlings were scored for three different morphological traits: Lesions, asymmetrical rosettes, and organ defects. The data represent two biological replicates (two replicates, n = 144 for *ago1* mutants, and n = 216 for Col-0 WT, *p<0.05, ttest). (**B**) Hypocotyl mean length and variance differ between wild-type and *ago1*-mutant seedlings. Hypocotyl length was measured for seven-day old, dark-grown seedlings. *ago1-27* mutant seedlings showed greater variance than Col-0 wild-type seedling in hypocotyl length (Levene’s test, p< 1.0E-03; n = 475 for *ago1-27*, n = 486 for Col-0 WT). Inset: boxplots of hypocotyl length means. Y-axis represents hypocotyl length (mm), **p< 1.0E-15, Mann-Whitney Wilcoxon test.

### *AGO1* buffers genetic variation independent of HSP90

We next tested whether *AGO1* perturbation could reveal cryptic genetic variation and whether *AGO1*-dependent loci overlapped with those buffered by *HSP90*. To do so, we introgressed the hypomorphic *ago1-27* allele into Col-0 lines with single chromosome substitutions from another, genetically divergent *Arabidopsis* strain, Landsberg *erecta* (L*er)* (STAIRS, STepped Aligned Inbred Recombinant Strains, **Figure 2A, Supplemental Table 2**). STAIRS lines have been generated for chromosomes 1, 3 and 5 (Koumproglou *et al*. 2002). Since *AGO1* is located on chromosome 1, we excluded these STAIRS lines from our analysis. For chromosomes 3 and 5, we selected two STAIRS lines each (chr3; N9448 and N9459, chr5; N9472 and N9501). The introgressed lines were genotyped to confirm the integrity of the respective L*er* segments.

**Figure 2.**
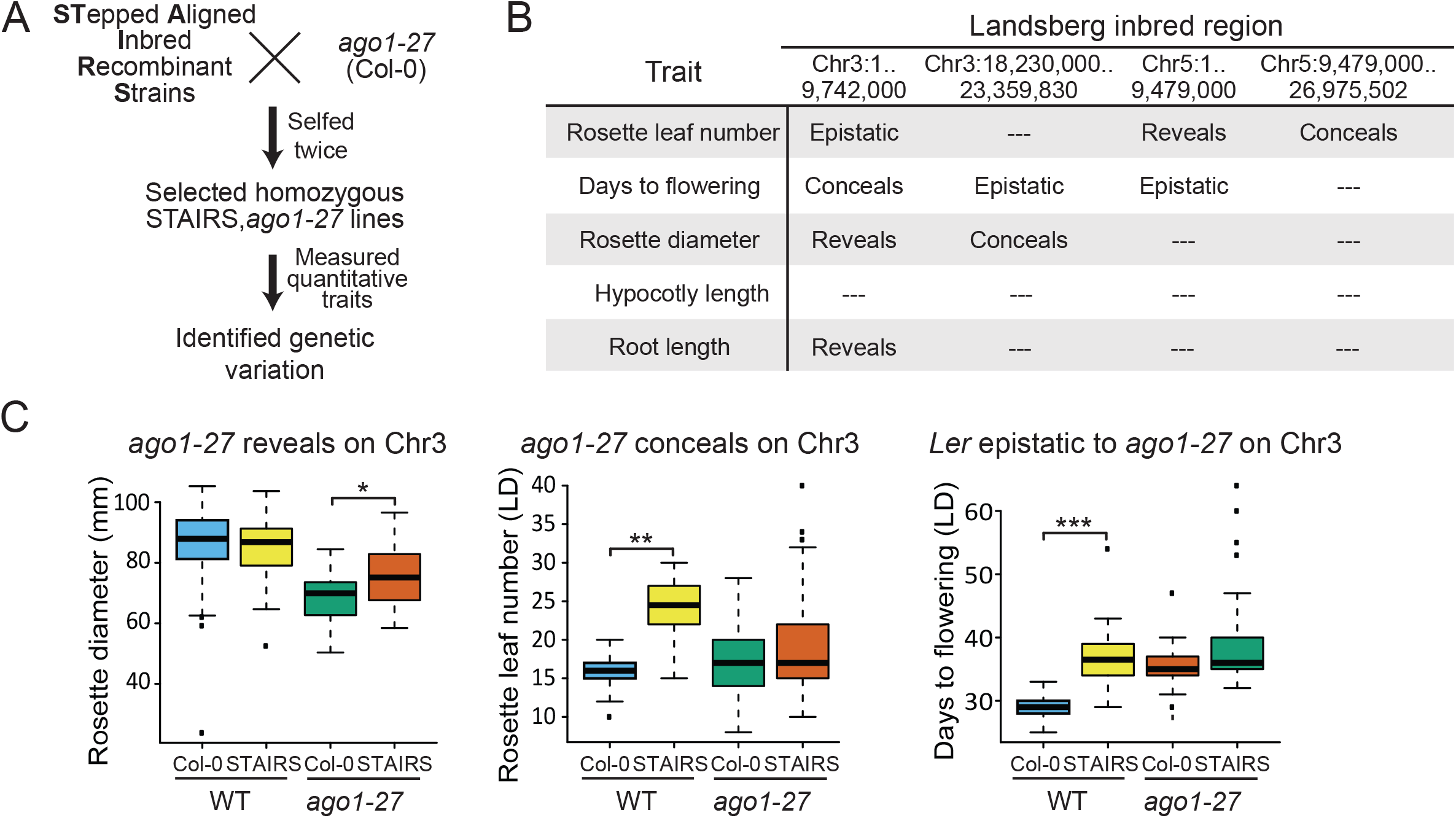
Perturbation of AGO1 buffers genetic variation. (**A**) Experimental design to examine the phenotypic consequences of genetic variation within the Stepped Aligned Inbred Recombinant Strains (STAIRS) in the context of the *ago1-27* mutation. (**B**) Summary of examined quantitative traits with evidence for *AGO1*-dependent or L*er*-dependent variation in each tested STAIRS line. AGO1 perturbation reveals a cryptic genetic variant if this variant’s contribution to a quantitative trait can be detected only in an *ago1*-mutant background. AGO1 perturbation conceals a genetic variant if this variant’s contribution to a quantitative trait can no longer be detected in an *ago1*-mutant background. Genetic variation in the respective L*er* segments can epistatically interact (*i*.*e*., mask) the phenotypic differences observed between Col-0 wild-type seedlings and *ago1-27* mutant seedlings in the Col-0 background. For STAIRS line N9472, 78 seedlings were measured for hypocotyl length in the dark, for STAIRS lines N9448, N9459, and N9501 100 seedlings were measured for this trait. At least 32 plants were measured for all other traits. See Supplemental Tables 2 and 3 for traits values and assessment of significance. (**C**) Two examples of *AGO1*-dependent and one example of L*er*-dependent genetic variation are shown for three different traits. Blue, Col-0 WT; Yellow, STAIRS; Red, *ago1-27* in the Col-0 background; Green, *ago1-27* in a STAIRS background.

We measured hypocotyl and root length, rosette diameter, and the closely correlated traits days to flowering and rosette leaf number across many individual plants per line using a randomized experimental design (**Supplemental Table 2**). We selected these traits because they are readily measurable and show evidence of HSP90-buffered variation in our previous studies of Col-0/L*er* mapping lines (Sangster et al. 2008a; Sangster et al. 2008b). Specifically, three previously described *HSP90*-dependent loci within the L*er* segments of the tested STAIRS lines affect the traits measured here (Sangster *et al*. 2008b; a).

*AGO1* perturbation may alter the contribution of a cryptic genetic variant to a quantitative trait in two ways: first, *AGO1* perturbation may reveal a genetic variant by increasing its contribution to a trait; or second, *AGO1* perturbation may conceal a genetic variant by increasing the relative contribution of others. Indeed, the phenomenon of revealing and concealing genetic variation has been previously observed for HSP90 perturbation across many traits in *Arabidopsis* recombinant inbred lines (Sangster *et al*. 2008a). In addition, genetic variation in the respective L*er* segments may mask the phenotypic differences observed between Col-0 wild-type and the *ago1-27* mutant that was generated in the Col-0 background (*i*.*e*., L*er* segments may epistatically interact with *ago1-27*). We observed all three scenarios of epistasis (**Figure 2B, C**, Supplemental Figures 2,**3**). Despite the strong evidence that HSP90 facilitates AGO1 function in many organisms, including plants, no overlap of *AGO1*-dependent loci with HSP90-dependent loci was observed.

### Perturbation of *AGO1* uncouples flowering time and rosette leaf number in a background-specific manner

One AGO1-buffered locus showed dramatically different effects on the two closely correlated traits days to flowering and rosette leaf number (**Figure 3A-C**). *Arabidopsis* plants develop about one rosette leaf per day until flowering is initiated. On average, Col-0 wild-type plants initiated flowering ~5 days later and have ~5 more leaves than the STAIRS line 9472 that carries a L*er* segment on chromosome 5 (coordinates 1 – 9,479,000bp). This result was expected because the L*er* segment in this STAIRS line encompasses the *FLOWERING LOCUS C (FLC)* gene (**Figure 3D**) which is non-functional in the L*er* strain (Michaels *et al*. 2003; Liu *et al*. 2004). *FLC* is a strong repressor of flowering (Whittaker and Dean 2017). In the Col-0 background, *ago1-27* plants initiated flowering ~9 days later and have ~2 more leaves, albeit the traits were less tightly correlated than in wild type **(Figure 3C**, compare blue and green dots). In stark contrast, in the STAIRS 9472 background, *ago1-27* plants showed no change in the number of days to flowering; however, these plants showed dramatically fewer leaves at the onset of flowering, developing on average only ~4 leaves. In fact, the severity of the rosette leaf number phenotype of STAIRS9472;*ago1-27* was comparable to that observed in loss-of-function early flowering mutants (Pouteau *et al*. 2004; Undurraga *et al*. 2012). In short, *AGO1* perturbation in the STAIRS line specifically affected the trait rosette leaf number while not affecting the trait days to flowering.

**Figure 3.**
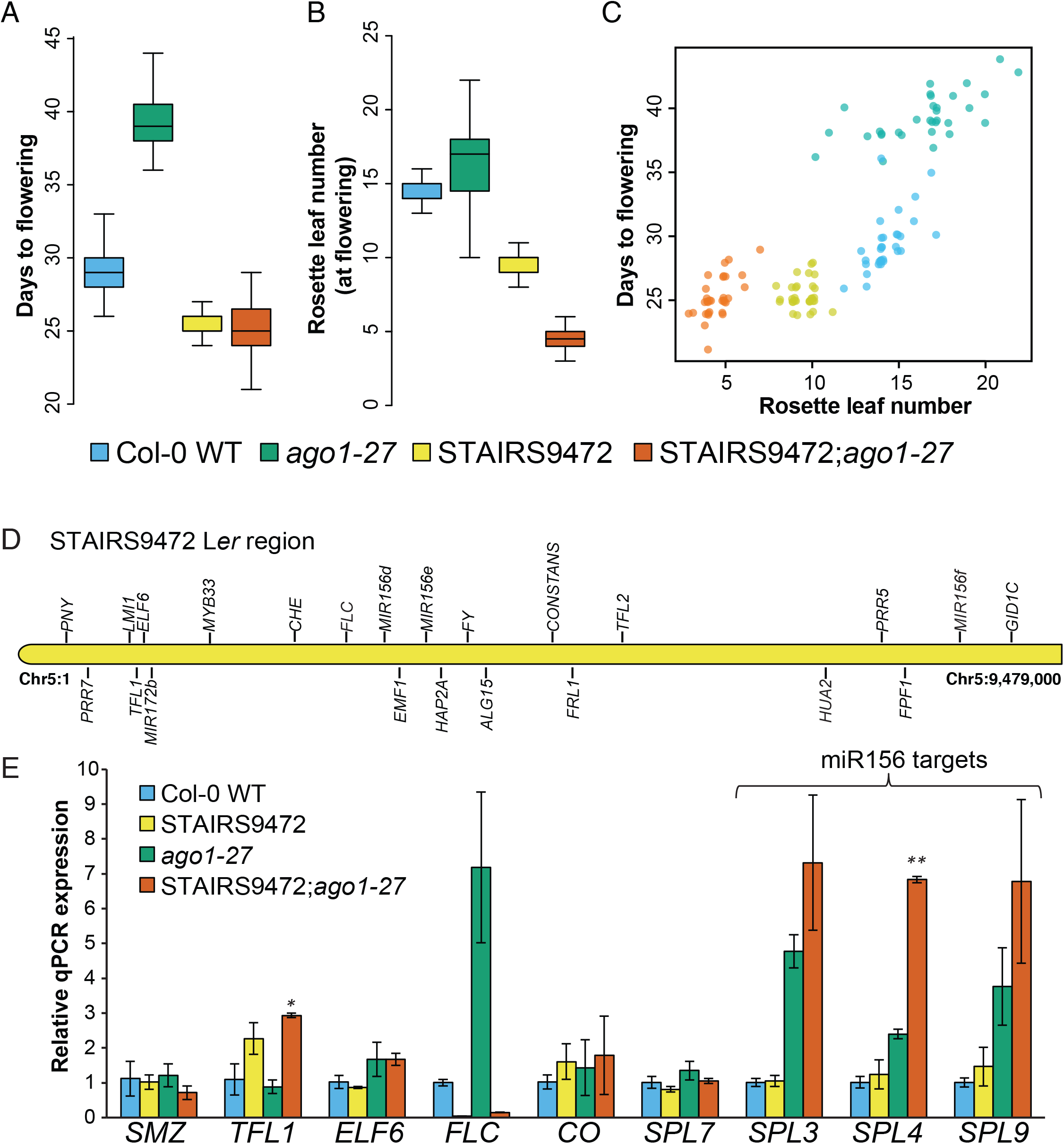
AGO1 perturbationuncouples the traits days to flowering and rosette leaf number. Plants for Col-0 WT, STAIRS9472, Col-0 *ago1-27* and STAIRS9472; *ago1-27* were grown in a random block design in long days, n = 30-36. Days to flowering were recorded and rosette leaf numbers at the onset of flowering were counted. Blue, Col-0 WT; Yellow, STAIRS9472; Red, *ago1-27* in the Col-0 background; Green, *ago1-27* in the STAIRS9472 background (**A**) Days to flowering. The *ago-1* mutant flowered ~9 days later than Col-0 WT (p=5.51E-12, Mann-Whitney Wilcoxon test); no significant difference was found between STAIRS9472 and the *ago-1* mutant in the STAIRS9472 background (p=0.4714, Mann-Whitney Wilcoxon test). The L*er* introgression in STAIRS9472 was epistatic to *ago1-27* *p<0.0001, Mann-Whitney Wilcoxon test. (**B**) Rosette leaf number. Col-0 WT plants showed fewer leaves than *ago1-27* mutant plants, consistent with the mutant’s late flowering phenotype. In the STAIRS 9472 background, *ago1-27* mutant plants showed no change in the number of days to flowering; however, these plants developed significantly fewer leaves (p=3.45E-12, Mann-Whitney Wilcoxon test). (**C**) Scatter plot of the two measured traits days to flowering and rosette leaf number in the four tested genotypes. The traits were less well correlated in the *ago-1* mutant in the Col-0 background (compare blue and green dots); however, the normally tight correlation was fully lost in the STAIRS background (compare red and yellow dots). (**D**) Known flowering time genes are residing within the L*er* chromosome 5 region of the STAIRS9472 line. (**E**) Quantitative PCR measurements for candidate gene expression. *TFL1* and *SPL4* were significantly increased in expression in the STAIRS9472;*ago1-27* background. 14-day old plant tissue was collected at ZT16 (Zeitgeber 16; 16 hours after dawn). Mean expression data represent two biological replicates, each with three technical replicates. Standard error is indicated. (* = p < 0.05, ** = p < 0.005, T-test).

### The close correlation of the traits days to flowering and rosette leaf number traits relies on *FLC* in the Col-0 background

The L*er* fragment in STAIRS9472 encompasses several known flowering time genes, including *FLC* which delays flowering by repressing the gene *FLOWERING LOCUS T (FT). FLC* expression is repressed when plants are exposed to cold temperatures for a prolonged period of time (*i*.*e*., vernalization or winter period), allowing *FT* expression and onset of flowering (Andrés and Coupland 2012; Whittaker and Dean 2017). Genetic variation in *FLC* and in *FRIGIDA (FRI)*, a positive regulator of *FLC*, accounts for the vast majority of differences in flowering time across *Arabidopsis* strains (Shindo *et al*. 2005; Kim *et al*. 2009; Bloomer and Dean 2017). Many *Arabidopsis* strains do not require vernalization to initiate flowering because they carry *FLC* mutations, as is the case for L*er*, or *FRI* mutations, as is the case for Col-0. The STAIRS9472 line carries the non-functional L*er FLC* allele.

We wondered if the lack of functional *FLC* in STAIRS9472;*ago1-27* contributed to its unusual phenotype. To test this possibility, we examined the consequences of repressing *FLC* through vernalization for both flowering time traits in Col-0 wild-type, STAIRS9472, Col-0 *ago1-27* and STAIRS9472;*ago1-27* plants (**Supplemental Figure 3**). Vernalization did not erase the difference in rosette leaf number between Col-0 *ago1-27* and STAIRS9472;*ago1-27* plants, with the latter still showing significantly fewer leaves (p = 5.704e^−12^ Wilcoxon test). However, vernalization uncoupled both flowering time traits in an *AGO1*-dependent manner in the Col-0 background. Although vernalized *ago1-27* plants initiated flowering ~5 days later than vernalized Col-0 wild-type plants, they had ~4 fewer leaves rather than more leaves. We conclude that the close association of days to flowering and rosette leaf number in the Col-0 background requires the presence of functional *FLC* and *AGO1*. Perturbation of *AGO1* alone diminished the close correlation between both traits but did not reverse it. L*er* and other natural *FLC* mutants must have rewired flowering time pathways such that the traits days to flowering and rosette leaf number remain closely correlated in the absence of functional *FLC*.

### *MIR156* polymorphisms are unlikely to cause *AGO1*-dependent phenotype

To identify the causal polymorphism(s) underlying the *AGO1*-dependent STAIRS9472 phenotype, we examined other genes within the L*er* segment with functions in flowering time (Song *et al*. 2013, 2015; Spanudakis and Jackson 2014) and candidate polymorphisms between Col-0 and L*er* (Nordborg *et al*. 2005; Borevitz *et al*. 2007; Ossowski *et al*. 2008) (**Figure 3D**). We measured expression of these candidate genes among the four genotypes Col-0 wild-type, STAIRS9472, Col-0 *ago1-27* and STAIRS9472;*ago1-27*; for the three *MIR156* genes (e, d, f), and *MIR172b*, we measured expression of major target genes (Ji *et al*. 2011). As expected, *FLC* expression was barely detectable in STAIRS9472 and STAIRS9472; *ago1-27* plants (**Figure 3D**), consistent with the known disruption of *FLC* in L*er* (Michaels *et al*. 2003; Liu *et al*. 2004). *FLC* expression increased in Col-0 *ago1-27* plants relative to Col-0 wild-type plants, consistent with the late flowering phenotype of the former genotype. As a general trend, target genes of *MIR156* increased in expression in the STAIRS *ago1-27* background compared to target gene expression in the Col-0 *ago1-27* background, suggesting that *MIR156* may be less functional in the STAIRS line. *MIR156* represses the expression of several *SQUAMOSA PROMOTER BINDING LIKE* (SPL) transcription factors (miR156-SPL module, **Figures 3E, 5D**) that regulate flowering by activating and repressing other transcription factors and miRNAs (Aukerman and Sakai 2003; Yamaguchi *et al*. 2009; Wu *et al*. 2009). Overexpression of *MIR156* leads to delayed onset of flowering with many more rosette leaves (Wu *et al*. 2009; Xu *et al*. 2016), suggesting that less functional *MIR156* may diminish rosette leaf number.

We searched for L*er*-specific polymorphisms in the *MIR156* genes in available genome assemblies and found a predicted single nucleotide polymorphism (SNP) within the loop of *MIR156f*. Resequencing of all three *MIR156* genes confirmed this SNP and identified an additional deletion of 14 nucleotides near the base of the stem loop. As the *MIR156* genes are highly conserved in the plant kingdom (Cuperus *et al*. 2011; Luo *et al*. 2013), we examined their natural variation among other *Arabidopsis* strains, sequencing an additional 55 strains. Of all sequenced strains, 42 carried the L*er*-specific C-to-T SNP, one carried a C-to-G SNP, and 32 carried the 14-nt deletion **(Supplemental Figure 2**). The presence or absence of the deletion was highly correlated with the presence or absence of the SNP (R^2^ = 0.3506, p = 0.0007, Pearson correlation test). To address whether either one or both L*er*-specific *MIR156f* polymorphisms affect rosette leaf number, we tested for association with this trait across these accessions (phenotypic data from (Lempe *et al*. 2005)). No association was found. Although this result did not rule out the *MIR156f* polymorphisms as the causative *AGO1*-dependent alleles, it made it less likely that these polymorphisms would explain the unusual STAIRS9472;*ago1-27* phenotype.

### Identifying the *AGO-1* dependent L*er*-specific polymorphism with a bulk segregant analysis

To identify the L*er*-specific variant(s) causing the observed trait uncoupling in STAIRS9472; *ago1-27* plants, we used a classic bulk segregant analysis followed by high-throughput sequencing (Cuperus *et al*. 2010; Sun and Schneeberger 2015). To generate a segregating population for the tested alleles, we crossed STAIRS9472 with *ago1-27*, and allowed for selfing to generate F_2_ seeds. F_2_ plants were measured for days to flowering and the number of rosette leaves at this point (**Figure 4A**). From this F_2_ experiment, we pooled plants based on phenotype, defining the STAIRS9472;*ago1-27* phenotype as plants with 6 or fewer rosette leaves (**Figures 3A-C, 4A)**, and isolated their DNA. We combined equal DNA amounts for 100 plants with the *AGO1*-dependent STAIRS9472 phenotype and 100 plants with higher numbers of rosette leaves. Using short-read sequencing, we aligned reads to the relevant chromosome 5 segment using SHOREmap (Sun and Schneeberger 2015), relying on the many known polymorphisms between L*er* and Col-0 to distinguish L*er*- and Col-0-specific reads. If successful, bulk segregant analysis will show increasing enrichment of homozygosity near the causal locus, with the causal locus at the center of a peak region (Salathia *et al*. 2007; Schneeberger *et al*. 2009; Cuperus *et al*. 2010; Sun and Schneeberger 2015). This mapping approach works best if variation at a single locus causes a segregating phenotype, and if phenotypes can be clearly distinguished from each another in order to pool samples with high confidence.

**Figure 4.**
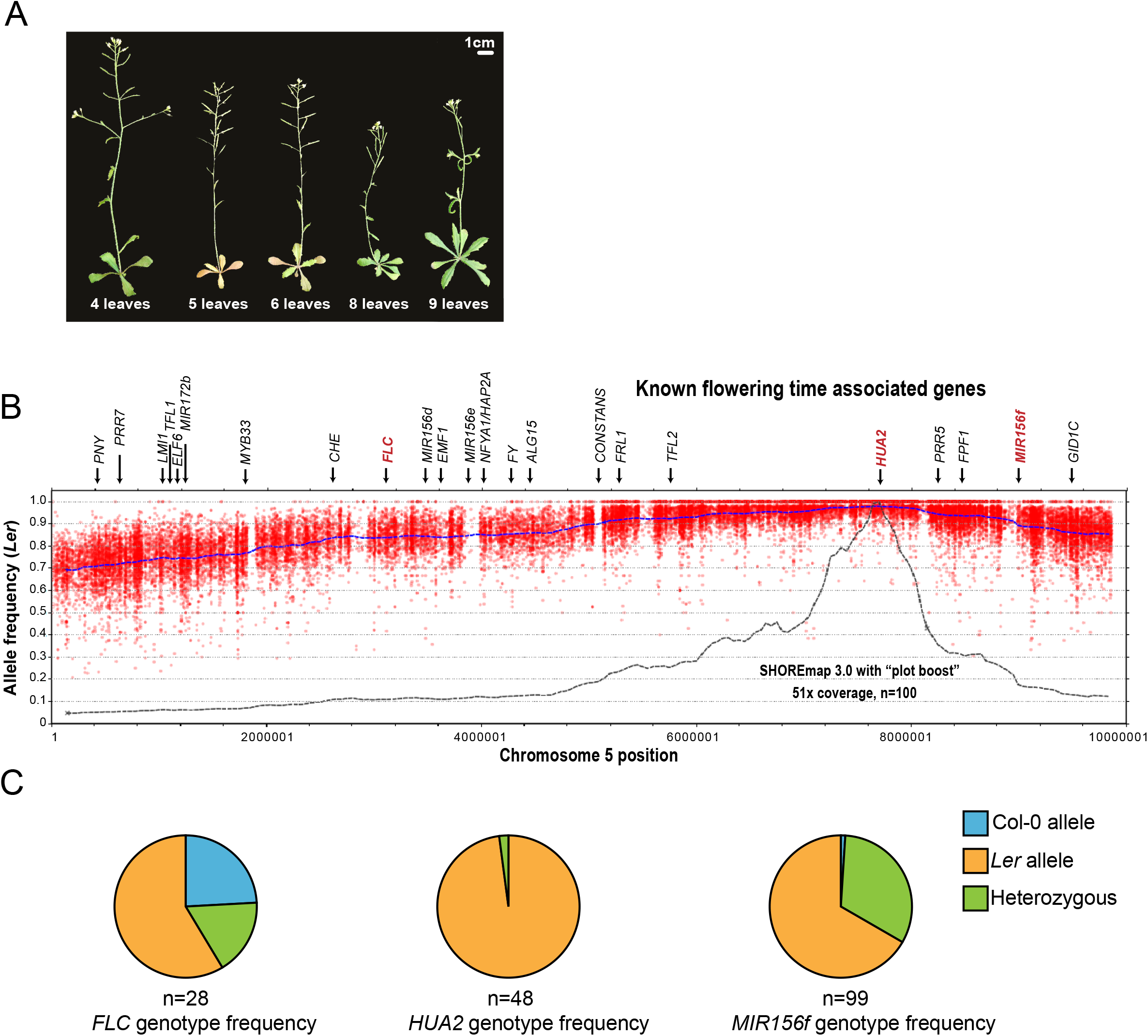
Bulk segregant analysis identifies the non-functional L*er hua2* allele as a candidate *AGO1*-dependent locus. (**A**) F_2_ plants from *ago1-27* x STAIRS9472 cross were grown in long days, phenotypes were recorded, and plants were genotyped for the *ago1-27* allele. For bulk segregant analysis, we selected plants that were homozygous for the *ago1-27* mutation and flowered with six or fewer leaves (n=100), resembling the *AGO1*-dependent STAIRS9472 phenotype. Representative F_2_ *ago1-27* plants at flowering are shown. Scale bar = 1cm. (**B**) Bulk segregant analysis. Red dots represent L*er* allele frequencies on chromosome 5 (bp, x-axis). Allele frequencies (y-axis) were estimated as the fraction of reads supporting a L*er* allele divided by the number of reads mapping to that locus. Dashed blue line represents sliding window-based allele frequencies as estimated by SHOREmap. Dashed black line represents window-based plot boost as estimated by SHOREmap. The L*er hua2-5* allele emerged as the candidate *AGO1*-dependent locus because L*er* allele frequencies were highest at this locus compared with other regions on chromosome 5. (**D**) F_2_ plants homozygous for the *ago1-27* mutation with six or fewer leaves at flowering were PCR genotyped for alleles at *FLC, HUA2*, and *MIR156f* loci. The near perfect enrichment of L*er hua2-5* allele validates the result of our bulk segregant analysis.

Although our phenotype of interest was quantitative in nature, *i*.*e*., a range of leaf numbers rather than an absence or presence of a feature, we observed a skew towards L*er* alleles on chromosome 5 with a SHOREmap peak region at chr5:7,600,000 to chr5:7,800,000 (**Figure 4B**). Of the known flowering time-associated genes, only one fell in this peak region, *HUA2* (AT5G23150). Some L*er* backgrounds, including the STAIRS9742 line, carry a premature stop codon mutation in *HUA2*, likely disrupting function (Chen and Meyerowitz 1999; Doyle *et al*. 2005; Zapata *et al*. 2016). *HUA2* function is less well characterized than that of other flowering time genes; however, *hua2* mutants in a Col-0 background show reduced *FLC* levels and fewer rosette leaves at onset of flowering (Doyle *et al*. 2005). *MIR156f* did not reside in the peak region, consistent with the previously described lack of genotype-phenotype association (**Figure 4B, C**).

To confirm that loss of functional *HUA2* was responsible for the *AGO1*-dependent phenotype in STAIRS9472, we used a Col-0-derived *hua2* mutant allele, *hua2-4*, and generated a double mutant *hua2-4;ago1-27* in the Col-0 background. We predicted that this homozygous double mutant would exhibit the uncoupling of days to flowering and rosette leaf number traits observed in the STAIRS9472; *ago1-27* line. Using a segregating F_2_ population, we simultaneously measured days to flowering and rosette leaf number, and genotyped each plant (**Figure 5A-C**). The *hua2-4* single mutant plants and the *hua2-4;ago1-27* double mutant plants showed no significant difference in days to flowering but rosette leaf number was markedly reduced in double mutant plants, recapitulating our original finding with STAIRS9472;*ago1-27* plants. The observed uncoupling of these traits was independent of *FLC* which is not disrupted in the *hua2-4;ago1-27* double mutant. This result suggests that the L*er*-specific, non-functional *hua2* allele may compensate for the L*er*-specific *FLC* disruption, thereby maintaining the close association of days to flowering and rosette leaf number traits.

**Figure 5.**
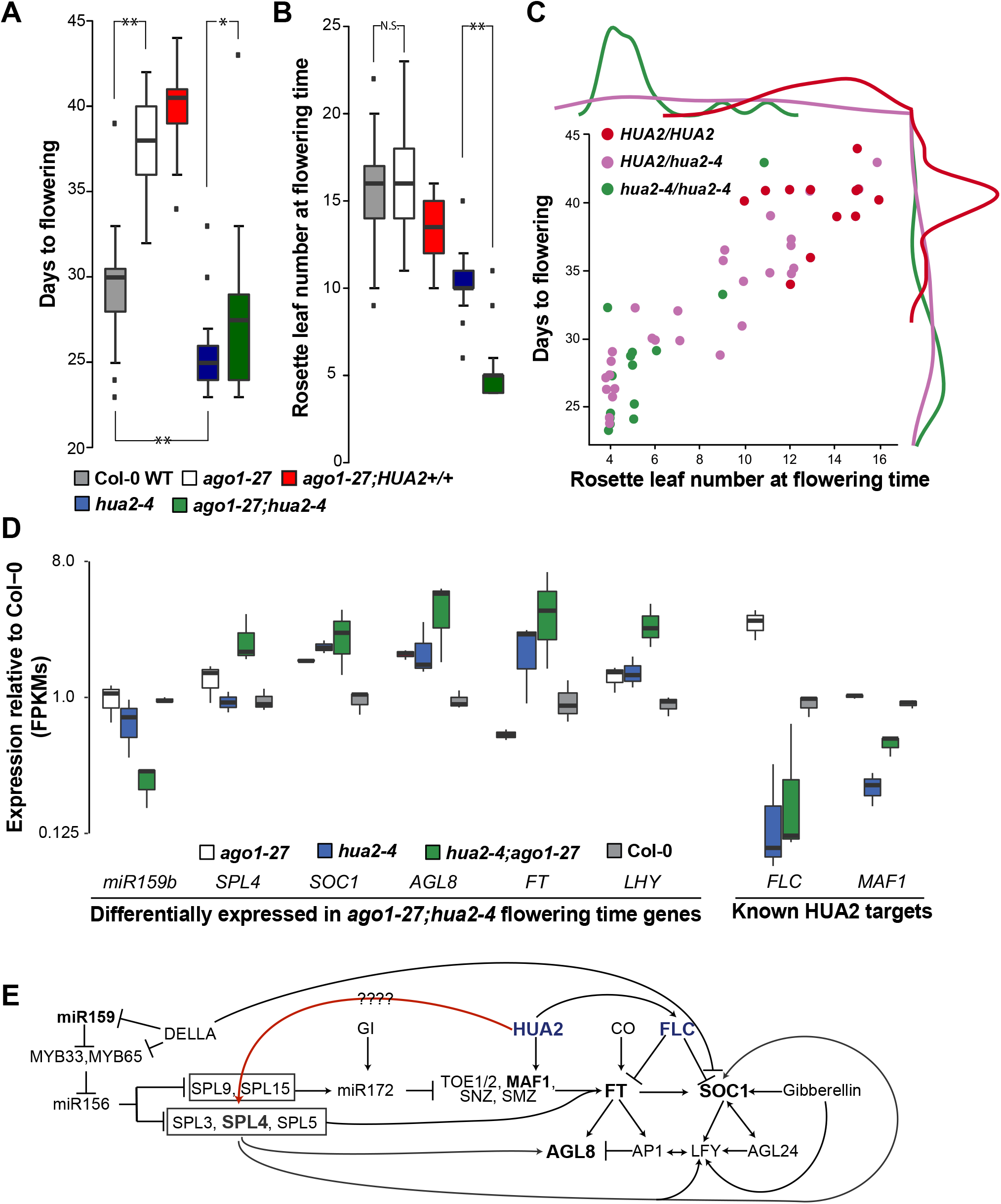
The *ago1-27*; *hua2-4* double mutant uncouples the traits days to flowering time and rosette leaf number in the Col-0 background. An F_2_ population segregating for the *ago1-27* and *hua2-4* mutant alleles was grown in long days. Days to flowering were recorded and rosette leaf numbers at the onset of flowering were counted. Grey, Col-0 WT; white *ago1-27* parent Red, *ago1-27;HUA2+/+* F2; Blue, *hua2-4* parent; Green, *ago1-27;hua2-4*. See Supplemental Table 5 for further details (**A**) Plants carrying a homozygous *ago1-27* allele flowered ~8.6 days later than Col-0 WT with ~16 leaves. Plants carrying a homozygous *hua2-4* allele initiated flowering ~4.5 days earlier than Col-0 WT. As observed for STAIRS9472;*ago1-27*, the *hua2-4* mutant allele was epistatic to *ago1-27*. *p<0.0283, **p<1.0E-06, Mann-Whitney Wilcoxon test. The double mutant *ago1-27;hua2-4* plants showed a similar mean value but greater trait variance. (**B**) The rosette leaf number phenotype of the double mutant *ago1-27;hua2-4* plants resembles that of the STAIRS9472;*ago1-27* line. *ago1-27;hua2-4* plants flower with 5 leaves on average. (*p<0.0283, **p<1.0E-06, Mann-Whitney Wilcoxon test, for both A and B) (**C**) Scatter plot with rosette leaf number on the x-axis and days to flowering on the y-axis. Data are shown for F_2_ plants that are homozygous for the *ago1-27* mutant allele and segregate for the *hua2-4* mutant allele. (*D*) Known flowering time genes with differential expression in the *ago1-27;hua2-4* double mutant as determined by RNA-seq. (**E**) Suggested intersection of miRNAs and HUA2 in flowering time pathways. Depicted in red is the proposed connection between *HUA2* and *SPL4* such that *SPL4* expression is regulated by both mir156 and HUA2. In bold, genes that are overexpressed in the double mutant *ago1-27,hua-2* relative to the single mutant *ago1-27*.

### *HUA2* effects on gene expression suggest *SPL4* as a likely *HUA2* target

To understand in more detail how *HUA2* affects the complex flowering gene network, we conducted RNA-seq experiments examining wild-type Col-0, single mutants *hua2-4* and *ago1-27* and *hua2-4;ago1-27* double mutant seedlings. As expected, *ago1-27* mutants and Col-0 wild-type showed differential expression of many miRNA target genes (**Supplemental Table 6**). The expression of the known *HUA2* targets *FLC* and *FLOWERING LOCUS M* (*FLM, MAF1*) was reduced in both the single *hua2-4* mutant and the *hua2-4;ago1-27* double mutant seedlings, excluding them as sources of the *AGO1*-dependent phenotype.

However, the comparison of *ago1-27* and *hua2-4;ago1-27* plants showed strong upregulation of *SPL4* expression in the latter (**Figure 5D**), consistent with our finding that *SPL4* was strongly upregulated STAIRS9472;*ago1-27* (**Figure 3E**). Other important flowering time genes were also differentially expressed in *hua2-4;ago1-27* double mutant plants, including the master regulator *FT, LATE ELONGATED HYPOCOTYL(LHY), SUPPRESSOR OF OVEREXPRESSION OF CONSTANS 1 (SOC1), AGAMOUS-LIKE 8 (AGL8, FRUITFUL)*, and *MIR159b* (**Figure 5D**). These genes interact in complex ways to control the transition to flowering (**Figure 5E**). Because *HUA2* is involved in mRNA processing and splicing (Chen and Meyerowitz 1999; Cheng *et al*. 2003; Janakirama 2013), we speculate that *SPL4* may be one of its targets. *SPL4* has three splice isoforms, and two of them (*SPL4-2, SPL4-3*) lack a miR156-binding site (Yang *et al*., 2012). Overexpression of *SPL4-1*, the splice form with the miR156 binding site, does not affect days to flowering but decreases rosette leaf number. In contrast, overexpression of *SLP4-2* or *SPL4-3* decreases days to flowering and reduces rosette leaf number (Yang *et al*. 2012). In a *hua2*;*ago1* double mutant background the balance of *SPL4* splice forms may be altered, which together with the absence of functional AGO1 disrupts the close correlation of days to flowering and rosette leaf number.

## Discussion

Here we show that AGO1, the principal player in miRNA-mediated control of gene expression in plants, buffers micro-environmental variation and maintains developmental stability in isogenic *Arabidopsis* seedlings. Compared to wild-type Col-0 plants, *ago1* mutant seedlings showed more lesions on cotyledons (Mason *et al*. 2016), more rosette symmetry defects and abnormal organs, and increased variation in hypocotyl length of dark-grown seedlings. Given the crucial role that miRNAs play in plant development, these results are not altogether surprising. MicroRNAs can impact developmental stability, *i*.*e*., the accuracy with which a given genotype produces a trait in a particular environment, in various ways (Hornstein and Shomron 2006; Voinnet 2009). For example, miRNAs can buffer gene expression noise as part of incoherent feedforward loops, in which a transcription factor will activate both the expression of a target gene X and a miRNA, with the latter repressing target gene X (Hornstein and Shomron 2006; Voinnet 2009). MicroRNAs enforce developmental patterning decisions through mutual exclusion and spatial or temporal restrictions in expression, *e*.*g*., by suppressing fate-associated transcription factors in neighboring cells or at a certain time in development (Hornstein and Shomron 2006; Voinnet 2009).

In plants, we previously reported increased variation for the same traits in isogenic seedlings upon perturbation of the chaperone HSP90 (Queitsch *et al*. 2002; Sangster *et al*. 2008b), consistent with the reported functional relationship of HSP90 and AGO1 (Iki *et al*. 2010, 2012; Iwasaki *et al*. 2010, 2015; Naruse *et al*. 2018). An exception was the peculiar environmentally-responsive lesions found on cotyledons in *ago1-27* seedlings (Mason *et al*. 2016). *HSP90* single mutants produce far fewer seedlings with lesions than *ago1-27* mutants, and double mutants show many more lesions than *ago1-27* single mutants, inconsistent with simple epistasis. Thus, we previously suggested that AGO1 is a major, but largely HSP90-independent, factor in providing environmental robustness to plants.

In addition to maintaining developmental stability, HSP90 buffers genetic variation in plants, fungi, and animals, including humans (Zabinsky *et al*. 2019). The hypothesized mechanism by which HSP90 overcomes the presence of genetic variation is the chaperone’s well-characterized function in protein folding and maturation (Sangster *et al*. 2004; Jarosz *et al*. 2010; Zabinsky *et al*. 2019). This hypothesis is supported by the reported differences among disease-associated protein variants chaperoned by HSP90 versus those chaperoned by HSP70 (Karras *et al*. 2017). Moreover, across thousands of humans, kinases that are HSP90 clients tend to carry more amino acid variants than non-client kinases, and these amino acid variants are predicted to be more damaging to protein folding (Lachowiec *et al*. 2015).

In contrast, it is harder to envision a simple, direct mechanism by which AGO1 overcomes the presence of genetic variation in either miRNAs or their targets, unless such buffering involves AGO1’s close relationship with HSP90 for the latter. Although we observed several instances in which AGO1 perturbation revealed and concealed genetic variation in the same traits in which we previously found HSP90-dependent variation (Sangster *et al*. 2008b), there was no overlap in the genetic loci buffered by AGO1 and HSP90. While this result was consistent with our study of the AGO1-dependent lesions (Mason *et al*. 2016), it raised anew the question as to how AGO1 may buffer genetic variation. In flies, proper expression of mir-9a, a miRNA acting on the transcription factor *Senseless*, buffers genomic variation (Cassidy *et al*. 2013). Reducing mir-9a regulation of *Senseless* leads to phenotypic variation in sensory cell fate in genetically diverse flies, with candidate causal variants in genes that belong to the *Senseless*-dependent proneural network governing sensory organ fate. In other words, in this case, AGO1-dependent buffering via mir-9a occurs at the network level, consistent with the mechanisms by which miRNAs buffer development stability and micro-environmental fluctuations.

To fully understand an instance of AGO1-dependent genetic variation, we focused on the uncoupling of the traits days to flowering and rosette leaf number in STAIRS9472. Both traits involve the mir156-SPL module and the key players *FLC* and *FRI* (**Fig. 5D**). We show that in the Col-0 background the coupling of these traits requires functional *FLC* and *AGO1*. In STAIRS9472, *FLC* is non-functional because the gene resides in the L*er*-introgression segment. Without *FLC*, how are days to flowering and rosette leaf number coupled in L*er*? Using bulk segregant analysis, we identified the non-functional *hua2* L*er*-allele as the likely causal AGO1-dependent polymorphism. Indeed, we were able to recapitulate the uncoupling phenotype in the *hua2-4; ago1-27* double mutant in the Col-0 background.

It is noteworthy that this non-functional *HUA2* allele arose only recently and likely in the laboratory (Zapata *et al*. 2016); there are several L*er* strains without this allele (Chen and Meyerowitz 1999; Doyle *et al*. 2005; Zapata *et al*. 2016). These strains and other *Arabidopsis* accessions with nonfunctional *FLC* genes must have acquired different polymorphisms to maintain the coupling of both traits. The inbreeding nature of *Arabidopsis* and the propagation of commonly used accessions like L*er* in controlled laboratory conditions readily allows fixation of such polymorphisms. However, there are prior reports that natural variation in *HUA2* can affect flowering time and plant morphology. The Sy-0 accession carries a gain-of function *HUA2* allele that enhances *FLC* expression leading to larger rosettes, in addition to suppressing *AGAMOUS* leading to indeterminate development of floral meristems (Wang *et al*. 2007). The Ws accession carries a 12bp-deletion in *HUA2*, possibly weakening *HUA2* function (Doyle *et al*. 2005).

Our expression analysis offered some clues as to how *HUA2* may facilitate the close coupling of days to flowering and rosette leaf number (**Figure 5D, E**). Comparing gene expression in *ago1-27* and *ago1-27; hua2-4* plants, we found that the mir156-SPL module gene *SPL4* was highly upregulated in the double mutant. *SPL4* is expressed in three splice isoforms (*SPL4-1, SPL4-2, SPL4-3*) with only one, *SPL4-1*, regulated by miR156 (Yang *et al*., 2012). Overexpression of *SPL4-1* in transgenic plants does not alter days to flowering but reduces rosette leaf number. In contrast, overexpression of *SLP4-2* or *SPL4-3* decreases both days to flowering and reduces rosette leaf number (Yang *et al*. 2012). Because *HUA2* functions in mRNA processing and splicing (Chen and Meyerowitz 1999; Cheng *et al*. 2003; Janakirama 2013), *SPL4* may be one of its targets. Non-functional *HUA2* may lead to increased presence of the *SPL4-1* splice form, which is exacerbated when mir156-dependent suppression of *SPL4-1* fails in the *ago1-27; hua2-4* double mutant, disrupting the close correlation of days to flowering and rosette leaf number (**Figure 5E**). Thus, similar to the scenario in flies (Cassidy *et al*. 2013), AGO1 appears to buffer genetic variation via microRNA-dependent network connections in plants. Disruption of the miRNA-dependent network path in *ago1*-mutants can reveal genetic variants such as the non-functional *HUA2* allele in other paths controlling the same trait (**Figure 5E**).

Taken together, our study holds several important lessons. First, AGO1 buffers phenotypic variation in isogenic seedlings and genetic variation in genetically divergent ones. Second, AGO1 does so independently of the chaperone HSP90 despite their close functional relationship, suggesting that epistasis is largely a first-order phenomenon, specific to two interacting loci. Indeed, this observed specificity of epistasis can extend to specific variants in pairwise interacting loci. We previously showed that HSP90 can buffer genetic variation in Ste12, a transcription factor that governs mating and invasion in yeast. However, HSP90-dependent variants in Ste12 are rare; they reside in only two positions that are close to one another and alter charge and DNA binding (Dorrity *et al*. 2018). This surprising specificity of epistatic interactions calls into question the utility of current large-scale efforts to understand the phenotypic contributions of epistasis by combining null mutants in human cells and in other organisms. Third, our results provide a cautionary tale in interpreting phenocopies. Mutants in AGO1 and HSP90 show highly similar phenotypes (Bohmert *et al*. 1998; Queitsch *et al*. 2002; Morel 2002; Vaucheret *et al*. 2004; Sangster *et al*. 2007; Mason *et al*. 2016), and yet the underlying mechanisms appear to differ, at least in part. Fourth, unlike HSP90, AGO1 suppresses the phenotypic consequences of genetic variation by enabling miRNA-dependent network paths rather than acting directly on variant-containing molecules, thereby extending the buffering concept. Fifth and last, we show that key pathways can involve different molecular players even in closely related strains of the same species. The uncoupling of highly correlated traits could be a useful tool for plant breeders who want to improve one trait without compromising another tightly coupled trait. Our study suggests miRNAs as good candidates for such targeted breeding and engineering efforts.

### Plant Materials and Growth Conditions

The following parental lines were used: Col-0, *ago1-27* in the Col-0 background, and STAIRS N9448, N9456, N9472, N9501(Morel 2002; Koumproglou *et al*. 2002). *ago1-27* plants were crossed into the STAIR lines and F_2_’s that carried the wild-type and *ago1-27* allele in both Col-0 and the STAIRS backgrounds were isolated. Selected F_2_’s and their progeny were used to perform the described experiments. For the hypocotyl and root length assays, the plants were grown on MS media containing 0.0005% MES hydrate, 0.004% vitamin solution, 3% phytoagar, and 1% sucrose.

### Genotyping of F_2_ plants

We used PCR to genotype the F_2_’s from each STAIRS – *ago1-27* cross. PCR conditions for *ago1-27* genotyping is as follows: 5’ at 94 °C, followed by 35 cycles at 30 s at 94 °C, 30 s at 55 °C, 1 min at 72 °C. PCR product was then digested at 37°C with Bsp1286I, which cuts wild-type sequence. PCR conditions for MIR156F genotyping is as follows: 2’ at 95°C, followed by 35 cycles at 30 s at 94°C, 50 s at 57°C, 40 s at 72°C.

### Hypocotyl and root length assays

Seeds from different genotypes were plated on agar plates (10 seeds/per plate, equally spaced). The plates were stacked in racks to ensure vertical position, wrapped in aluminum foil, and transferred to 4°C for five days to promote germination. They were then unwrapped, and exposed to light for two hours. After that, the plates were wrapped in aluminum foil again, to prevent further light exposure and were transferred to a 23°C tissue culture incubator for seven days. The plants were grown vertically. After that, the plates were taken out, and photographed. The photographs were used to measure the seedlings’ hypocotyls and roots using the ImageJ software (http://rsbweb.nih.gov/ij/).

### Early morphology traits analysis

Seeds from the different genotypes were plated on agar (36 seeds/per plate). The plates were wrapped in aluminum foil and transferred to 4°C for five days. Plates were unwrapped and transferred to long days (LD) in 23°C tissue culture incubator for 10 days. The plants were grown horizontally. The plates were rotated every day to prevent biases due to location in the incubator. On the 10^th^ day, the seedlings were scored for their morphological traits.

### Flowering time experiments

Seeds from different genotypes were embedded in 1ml of 0.1% agar, and then stratified for five days at 4°C. They were sown on soil in 36-pot trays. Flowering time was measured by scoring both the number of rosette leaves and days to flowering when the primary inflorescence of the plant had reached a height of 1cm. Flowering time experiments were performed in long days (LD, 16 hours of light, 8 hours of dark), at 23°C.

### Rosette diameter measurements

The diameter of the rosette was measured on the day that the primary inflorescence of the plant reached a height of 1cm.

### Vernalization treatment

Seeds were stratified for five days at 4°C and then sown on soil. They were allowed to grow for five days at 23°C in LD or short days (SD) conditions and then transferred to 4°C for forty days, according to recommendations from Sung *et al*., 2006 (Sung et al. 2006).

### Gene expression analysis

To determine the expression levels via qPCR, total RNA was isolated from the aerial parts of 14-day old plants at ZT16 using the SV Total Isolation System (Promega). RNA quality was determined using a Nanodrop and only high-quality samples (A260/A230 > 1.8 and A260/A280> 1.8) were used for subsequent qPCR experiments. To remove possible DNA contamination, RNA was treated with DNaseI (Ambion) for 60 minutes at 37°C. We used the Transcriptor First Strand cDNA Synthesis Kit (Roche) for cDNA synthesis. The qPCR primers were designed using the Universal Probe Library Assay Design Center tool (Roche), and Primer3 (Untergasser et al. 2012). Specific amplification was confirmed before conducting the qPCR experiments. The qPCRs experiments were carried out in 96-well plates with a LightCycler480 (Roche) using SYBR green. The following program was used for the amplification: pre-denaturation for 5 min at 95°C, followed by 35 cycles of denaturation for 15 s at 95°C, annealing for 20 s at 55°C, and elongation for 30 s at 72°C. All qPCR experiments were carried out with two biological replicates (independent samples harvested on different days), and with three technical replicates per sample.

RNA-seq samples were prepared similarly as for qPCR, and then using the Illumina stranded Tru-seq kit following the standard protocol. Samples were sequenced using the Nextseq550 platform. We used TopHat (v2.1.2) to align RNA-seq reads to the TAIR10 genome annotation (Trapnell *et al*. 2009), htseq-count (v0.12.4) to calculate counts per gene (Anders *et al*. 2015), using a minimum map quality of 10 and Cuffdiff (v2.2.1) to generate FPKMs (Trapnell *et al*. 2013), and DESeq2 to identify differentially expressed genes among genotypes (Love *et al*. 2014).

### Sequencing of miR156F, D and E in diverse *A. thaliana* strains

The genes *MIR156F, MIR156D* and *MIR156E* were amplified using the primers listed on **Supplemental Table 7**. PCR products were sequenced by the Sanger method. The sequences were aligned using T-coffee.

### Bulk segregant analysis - library preparation and sequencing

Approximately 400 F_2_ plants were sown, and leaf samples of equal size were collected from 100 plants that resembled the STAIRS9472;*ago1-27* phenotype (6 leaves or fewer at flowering) and 100 plants with a greater number of leaves. Individual plants were genotyped. In parallel, leaf samples were collected for all genotypes. DNA was extracted using CTAB extraction (Weigel and Glazebrook 2002) and quantified using the Qubit HS dsDNA assays. Libraries were quality checked on the Agilent 2100 bioanalyzer using a DNA 1000 chip (Agilent). Samples were pooled and libraries were generated using the Nextera sample kit according to the manufacturer’s instruction. DNA concentration of the amplified libraries was measured with the DNA 1000 kit as well as the DNA high-sensitivity kit for diluted libraries (both Agilent). Samples were sequenced on an Illumina Nextseq in a 75-bp paired-end run.

### Bulk segregant analysis – data analysis

Using the function SHORE import, raw reads were trimmed or discarded based on quality values with a cutoff Phred score of +38. After correcting the paired-end alignments with an expected insert size of 300 bp, we applied SHORE consensus to identify variation among mutants and reference. We applied SHOREmap using the included L*er*/Col-0 SNPs. Plot boost was applied to further define a mapping interval.

## Data availability

All RNA sequencing reads and genomic sequence used for the SHOREmap analysis are deposited in the NCBI Sequence Read Archive under the BioProject accession PRJNA836875.

## Acknowledgements

This work was supported by the National Human Genome Research Institute Interdisciplinary Training in Genomic Sciences (T32 training grant HG000035 to G.A.M.) and the National Institute of Health (New Innovator award DP2OD008371 and NIGMS 1R35GM139532-01 to C.Q.). We thank the editor and two anonymous reviewers for insightful comments and additional references that have improved this manuscript.

## Figure Legends

**Supplemental Figure 1**.Five phenotypes were measured for four STAIRS inbred lines, resulting in cases of revealing, concealing and epistatic phenotypic changes. The five phenotypes measured were days to flowering, rosette leaf number, rosette diameter, hypocotyl length and root length.

**Supplemental Figure 2**. (A) Landsberg, Col-0 and the inbred strain STAIRS9472 we sequenced across the MIR156F gene, supporting that there was both a SNP and a 14-nucleotide deletion in both the Landsberg and STAIRS9472 strains compared to Col-0. (B) Further sequencing across 55 A. thaliana strains. Of the sequenced strains,, 42 carried the L*er*-specific C-to-T SNP, one carried a C-to-G SNP, and 32 strains carried the 14-nt deletion.

**Supplemental Figure 3**. Seeds were stratified for five days at 4°C and then sown on soil. They were allowed to grow for five days at 23°C in LD or short days (SD) conditions and then transferred to 4°C for forty days to vernalize plants. (**A**) Rosette leaf number at flowering with vernalized plants in long day conditions. (**B**) Days to flowering with vernalized plants in long day conditions.

## Notes

### Competing Interest Statement

The authors have declared no competing interest.

### Summary of Updates

Minor changes to the manuscript, primarily in the introduction and discussion

