## Supplementary figures and images for "AGO1 and HSP90 buffer different genetic variants in *Arabidopsis thaliana*"

### Supplemental Figure 1

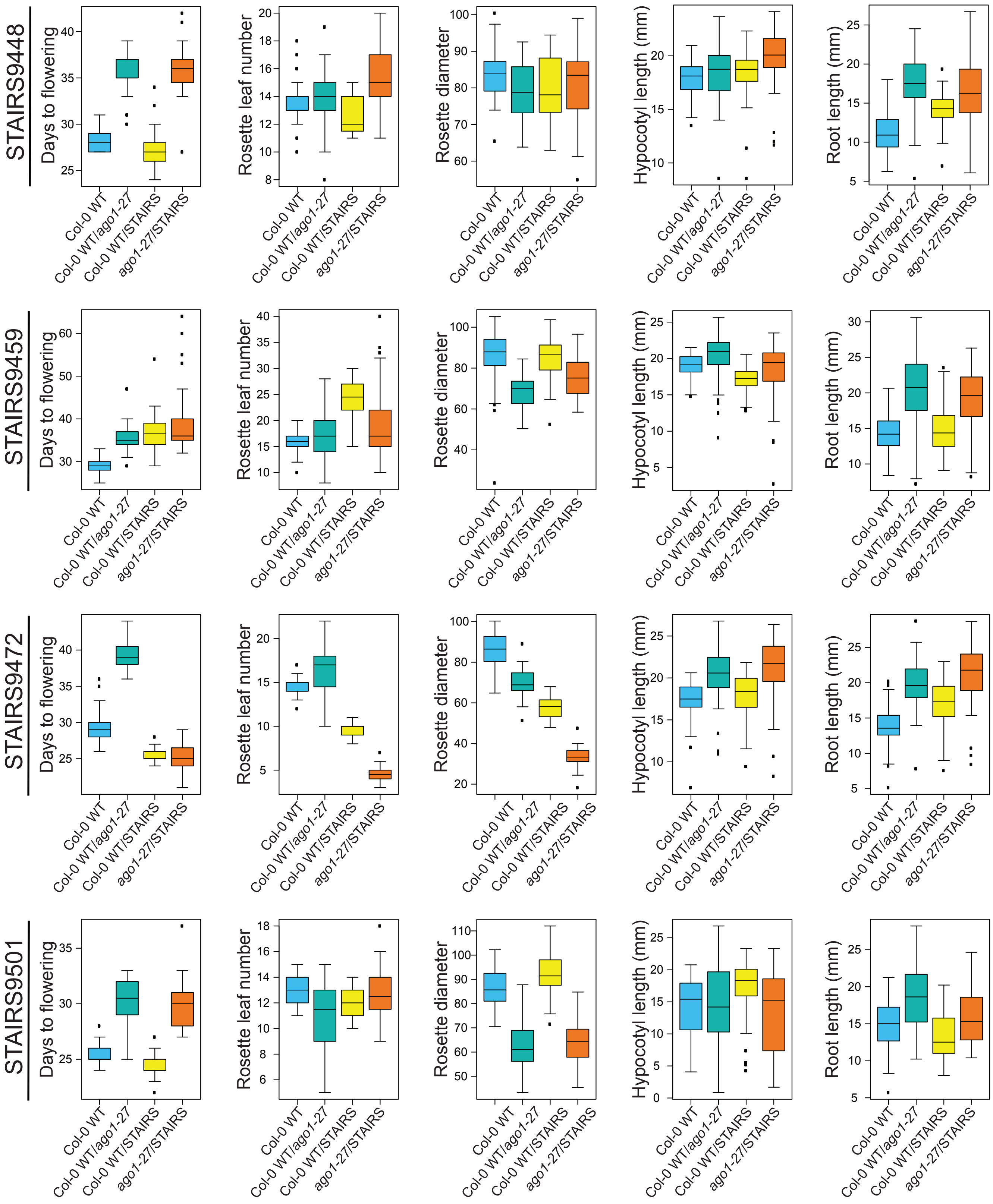

### Supplemental Figure 2

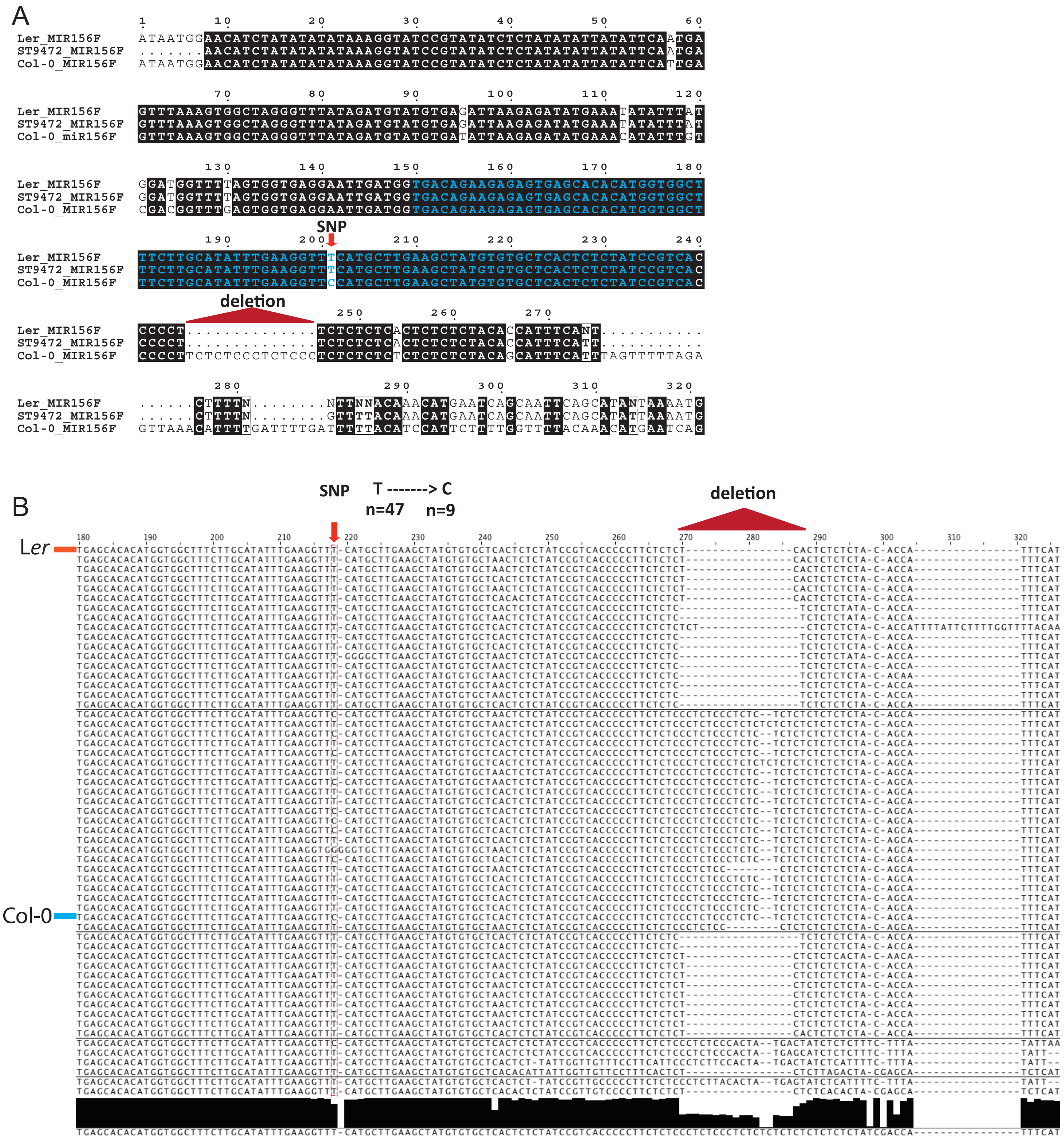

### Supplemental Figure 3

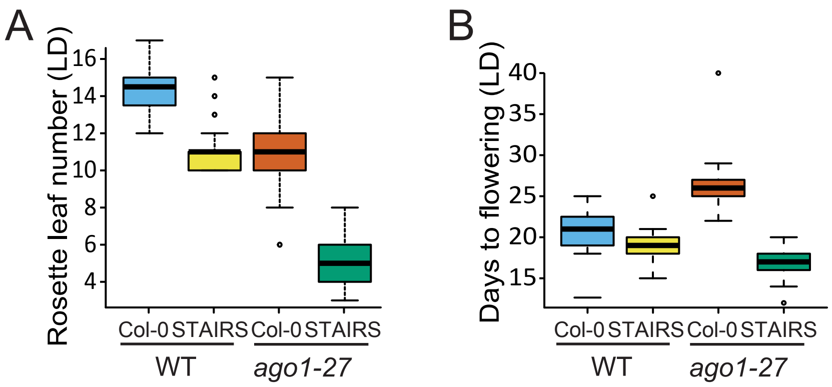
